# Deduplication Improves Cost-Efficiency and Yields of *De novo* Assembly and Binning of Shot-Gun Metagenomes in Microbiome Research

**DOI:** 10.1101/2022.10.12.512008

**Authors:** Zhiguo Zhang, Lu Zhang, Ze Zhao, Hui Wang, Feng Ju

## Abstract

Metagenomics has in the last decade greatly revolutionized the study of microbial communities. However, the presence of artificial duplicate reads mainly raised from the preparation of metagenomic DNA sequencing library and their impacts on metagenomic assembly and binning have never brought to the attention. Here, we explicitly investigated the effects of duplicate reads on metagenomic assembly and binning, based on analyses of four groups of representative metagenomes with distinct microbiome complexity. Our results showed that deduplication considerably increased the binning yields (by 3.5% to 80%) for most of the metagenomic datasets examined thanks to improved contig length and coverage profiling of metagenome-assembled contigs. Specifically, 411 versus 397, 331 versus 317, 104 versus 88 and 9 versus 5 metagenome-assembled genomes (MAGs) were recovered from MEGAHIT assemblies of bioreactor sludge, surface water, lake sediment, and forest soil metagenomes, respectively. Noticeably, deduplication reduced the computational costs of metagenomic assembly including elapsed time (by 9.0% to 29.9%) and maximum memory requirement (by 4.3% to 37.1%). Collectively, it is recommended to remove duplicate reads in metagenomic data before assembly and binning analyses, particularly for complex environmental samples, such as forest soils examined in this study.

**Importance:** Duplicated reads are usually considered as technical artefacts. Their presence in metagenomes would theoretically not only introduce bias in the quantitative analysis, but also result in mistakes in coverage profile, leading to negative effects or even failures on metagenomic assembly and binning, as the widely used metagenome assemblers and binners all need coverage information for graph partitioning and assembly binning, respectively. However, this issue was seldomly noticed and its impacts on the downstream key bioinformatic procedures (e.g., assembly and binning) still remained unclear. In this study, we comprehensively evaluated for the first time the impacts of duplicate reads on de novo assembly and binning of real metagenomic datasets by comparing assembly quality, binning yields and the requirements of computational resources with and without the removal of duplicate reads. It was revealed that deduplication considerably increased the binning yields and significantly reduced the computational costs including elapsed time and maximum memory requirement. The results provide empirical reference for more cost-efficient metagenomic analyses in microbiome research.

## Introduction

High-throughput DNA sequencing or next generation sequencing (NGS) technologies have revolutionarily evolved and continually commercialized over the past 20 years, which nowadays enable researchers to directly obtain genomic DNA sequences of whole microbial communities (i.e., metagenomes) at an unprecedently low sequencing cost (1). The reconstruction of microbial genomes from NGS data via metagenome assembly and binning enables systematic genomic and in-depth qualitative analyses ideally of largely uncultured microbes in environmental samples, not just those cultivable ones, offering novel insights into the microbial community functions and metabolic pathways in complex microbial systems (2,3). Since its first application, there has been spectacular successes in recovering thousands of metagenome-assembled genomes (MAGs) for uncultured taxa from various natural and engineered ecosystems (4–6), opening the gate to the intriguing microbiome sciences underlying our earth environment (e.g., biogeochemistry, eco-restoration, and bioremediation), bioeconomy (e.g., bioresource and bioenergy) and human systems (e.g., food and health) (7).

Duplicate reads of artificial origins, mainly result from sequencing two or more copies of the same DNA fragment amplified during the PCR amplification in library construction step, are a major technical concern in Illumina high-throughput metagenomic sequencing (8,9). Comparatively, natural duplicate (potentially from genomic DNA shearing at the same position in different molecules) frequency is expected to be extremely low, especially for complex environmental microbiomes lacking dominant species and/or with high alpha-diversity (10). The removal of artificial duplicates was of critical concern in whole genome sequencing, as duplicate reads can increase the potential biases on variant calling algorithms, in which the amplification-induced error of PCR duplicates may be misidentified as a true variant (11). In fact, failure to remove duplicate reads can also lead to incorrect conclusions in metagenomic analysis, for example, in the quantification of microbial taxa, genes, and metabolic pathways (12). Deduplication option was integrated in the standard metagenomic pipeline of MG-RAST which conducts read-based annotation analysis (13), while it has recently been included as a quality control step in metagenomic assembly and binning tools, such as ATLAS pipeline (14) and bhattlab_workflows (https://github.com/bhattlab/) which was adopted by the recent metagenomic research of gut microbiomes (15). Theoretically, the widely used metagenome assemblers and binners all need coverage information for graph partitioning and assembly binning, respectively (2,3), where duplicate reads might result in bias in computing coverage profile (e.g., uneven coverage), leading to negative effects or even failures on metagenomic assembly and binning of genomic information of microbial species. For instance, a better performance in the assembly of metagenomic sequencing data from amplification-free library was observed, comparing with that of amplification library containing much more duplicate reads (9). However, the actual impacts of duplicate reads in metagenomic data on the downstream assembly and binning, the two core bioinformatic procedures of today’s mainstream metagenomic methodology (2,3), remain elusive.

In this study, we comprehensively evaluated for the first time the impacts of duplicate reads on *de novo* assembly and binning of real metagenomic datasets by comparing assembly quality, binning yields and the requirements of computational resources with and without the removal of duplicate reads. To represent different environmental microbiomes, 36 metagenomes from four typical ecosystem habitats (i.e., bioreactor sludge, surface water, lake sediment, and forest soil) with different degrees of microbiome complexity were sampled and sequenced. We chose the widely-used metaSPAdes and MEGAHIT as assemblers and metabat2 as the binner to benchmark the evaluation process. MetaSPAdes is currently considered to be the best assembler for generating longer or better metagenome assemblies in general, while MEGAHIT is a faster-effective and lower-cost solution when computational resources are a limiting factor, according to previous assessment studies (16,17).

## Results

### Comparison of metagenomic assembly with and without deduplication

We firstly evaluated the microbiome complexity by analyzing metagenomic datasets of four representative ecosystem habitats, namely bioreactor sludge, surface water, lake sediment, and forest soil. The results confirmed that the forest soil metagenomes are the most complex among the examined metagenomic datasets, followed by lake sediment, surface water and bioreactor sludge metagenomes (Fig. 1). On average, an affordable and regular sequencing depth of 10Gbp of shot-gun metagenomes was estimated to cover 91.0%, 65.4%, 49.5% and 41.1% of the total microbiome diversity in bioreactor sludge, surface water, lake sediment, and forest soil samples, respectively. Deduplication removed 15.9% to 25.1% reads from clean data (raw data after quality filtering, see methods). Then, the clean data and deduplicated data of each sample in the four groups of metagenomic datasets were assembled individually using MEGAHIT and metaSPAdes. We used four popular metrics, i.e., Number of contigs>1kb, Number of contigs>10kb, Longest contig length, and N50 length, to comprehensively evaluate the assembly quality (Dataset S2). The assembly quality was significantly negatively correlated with microbiome complexity for both clean data and deduplicated data (Fig. S1). In general, bioreactor sludge metagenomes showed the highest assembly quality, followed by those of surface water, lake sediment and forest soil.

**Fig. 1.**
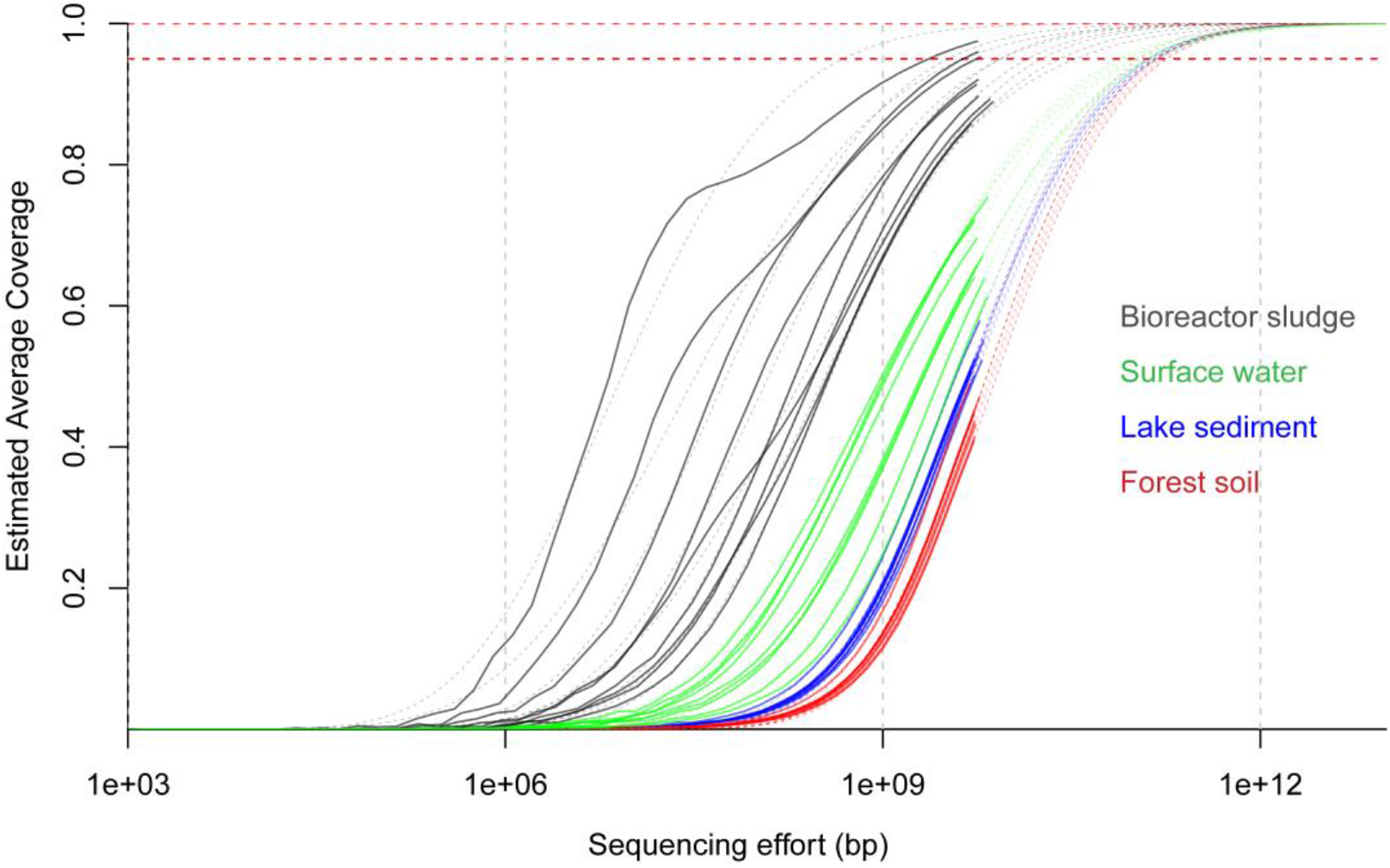
Nonpareil estimates of sequence coverage (redundancy) for the 36 metagenomes studied. Metagenomes are grouped according to their environmental niche, red, blue, green and black lines indicate forest soil, lake sediment, surface water and bioreactor metagenomes, respectively. Solid and dashed lines indicate real curve and model prediction, respectively. Sequencing effort is indicated in base pairs on a log scale and the estimated coverage achieved is shown as a fraction of 1.

For MEGAHIT assemblies, deduplication considerably improved the overall length of metagenome-assembled contigs, as the N50 length of the assemblies increased by 1.7% on average (Fig. 2a). However, deduplication decreased the total number of contig (>1000bp) by 2.9%, 3.2%, 5.1%, 7.9% on average for bioreactor sludge, surface water, lake sediment, and forest soil metagenomes (Fig. 2c). In forest soil metagenomes, which has the highest microbiome diversity in this study, the number of contigs with length > 10000 bp significantly increased by 17.6% after deduplication (wilcoxon signed-rank test, *P* = 0.004) (Fig. 2d). For metaSPAdes assemblies, Significant difference in N50 was not observed, except for the forest soil samples, for which N50 was slightly lower with deduplication (< 1% on average) (Fig. 2e). The number of contigs with length > 10000bp significantly (*P* = 0.006) decreased after deduplication in bioreactor sludge samples whereas increased in seven of nine forest soil metagenomes (Fig. 2h). In addition, the longest contig in 18 of 36 samples was 7.4% to 71.3% longer than those without deduplication (Fig. 2f). Interestingly, the total number of contigs (>1000bp) assembled from forest soil metagenomes slightly increased after deduplication (Fig. 2g), in contrast with MEGAHIT assemblies. In general, deduplication improved the length of metagenome-assembled contigs, but the total number of contig (>1000bp) was reduced by deduplication, particularly in bioreactor sludge, surface water and lake sediment metagenomes.

**Fig. 2.**
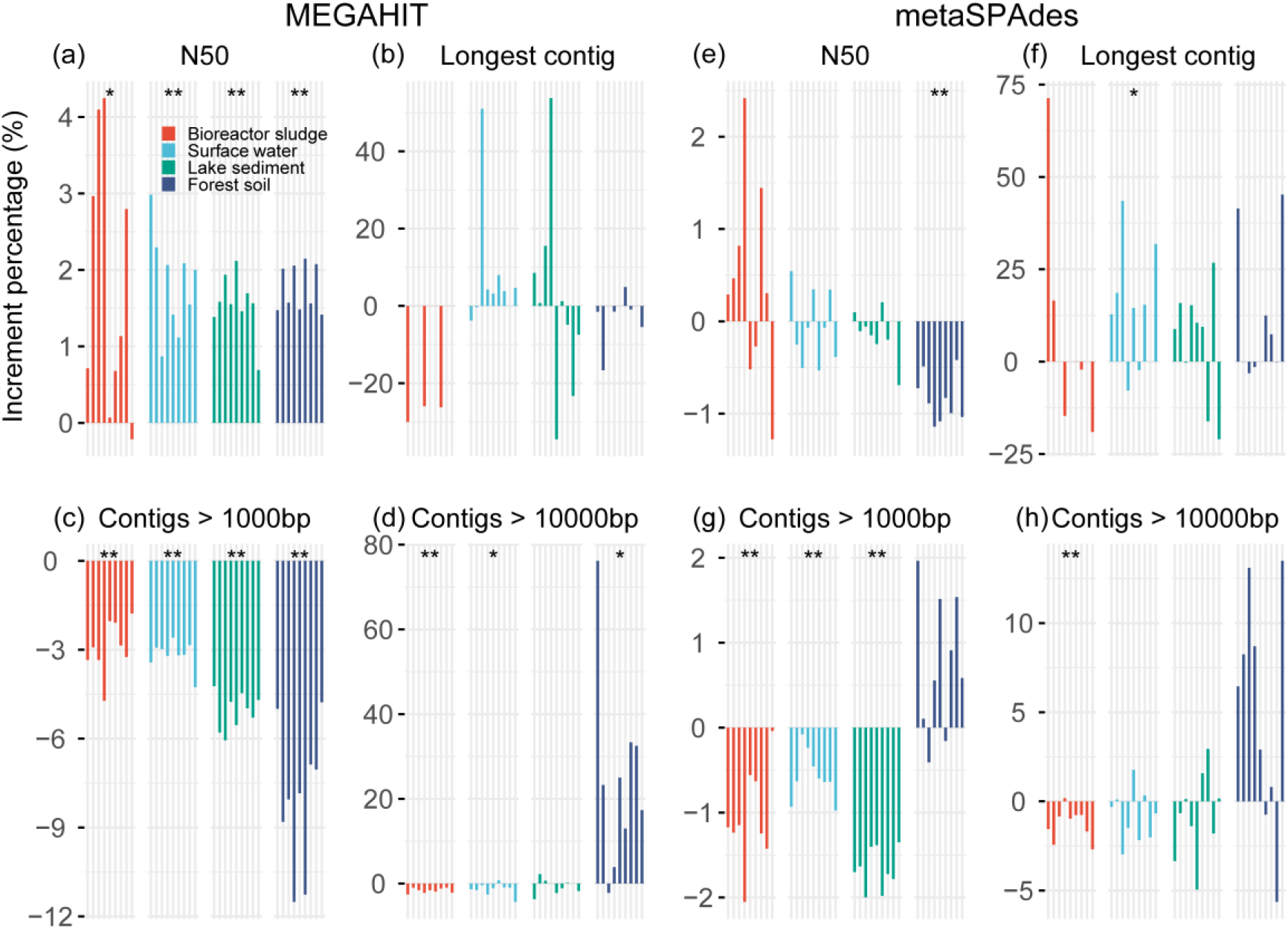
Comparison of assembly results between clean and deduplicated metagenomic data. Percentage ratio = (metric of deduplicated data - metric of clean data)/ metric of clean data. Significance was checked using wilcoxon signed-rank test. “*” and “**” indicate *P* < 0.05 and *P* < 0.01, respectively.

### Comparison of metagenomic binning with and without deduplication

Deduplication contributed to a better recovery of MAGs, as revealed by binning of both MEGAHIT and metaSPAdes assemblies (Fig. 3) using MetaBAT2. A total of 855 and 807 MAGs with over 50% quality (completeness-5× contamination) were recovered from 36 MEGAHIT assemblies of deduplicated data and clean data, respectively. For details, there were 411 versus 397, 331 versus 317, 104 versus 88 and 9 versus 5 MAGs from bioreactor sludge, surface water, lake sediment, and forest soil metagenomes, respectively (Fig. 3a). The results also indicated that, at the same level of 10Gbp sequencing depth (Dataset S1), fewer bins were recovered from metagenome datasets with higher microbiome complexity (Fig. 3a). Deduplication also contributed to the recovery of high-quality MAGs. For example, 11, 10, 13 and 2 more bins with completeness over 70% were recovered from deduplicated data of bioreactor sludge, surface water, lake sediment and forest soil metagenomes, respectively (Table 1). Furthermore, to compare the taxonomic diversity of MAGs recovered from clean data and deduplicated data, the MAGs were clustered using a 95% cut-off of whole-genome average nucleotide identity (ANI) to generate species-level representative MAGs. Results showed that the number of species-level MAGs recovered from deduplicated data was consistently higher than that of clean data for all four metagenomic datasets. In other words, there were 211 versus 206, 91 versus 90, 26 versus 25 and 5 versus 2 species-level representative MAGs recovered from bioreactor sludge, surface water, lake sediment, and forest soil metagenomes, respectively (Fig. 3a). When looking into their composition, it was found that most of MAGs were recovered from both deduplicated data and clean data, whereas more unique MAGs were recovered from deduplicated data (Fig. S2).

**Fig. 3.**
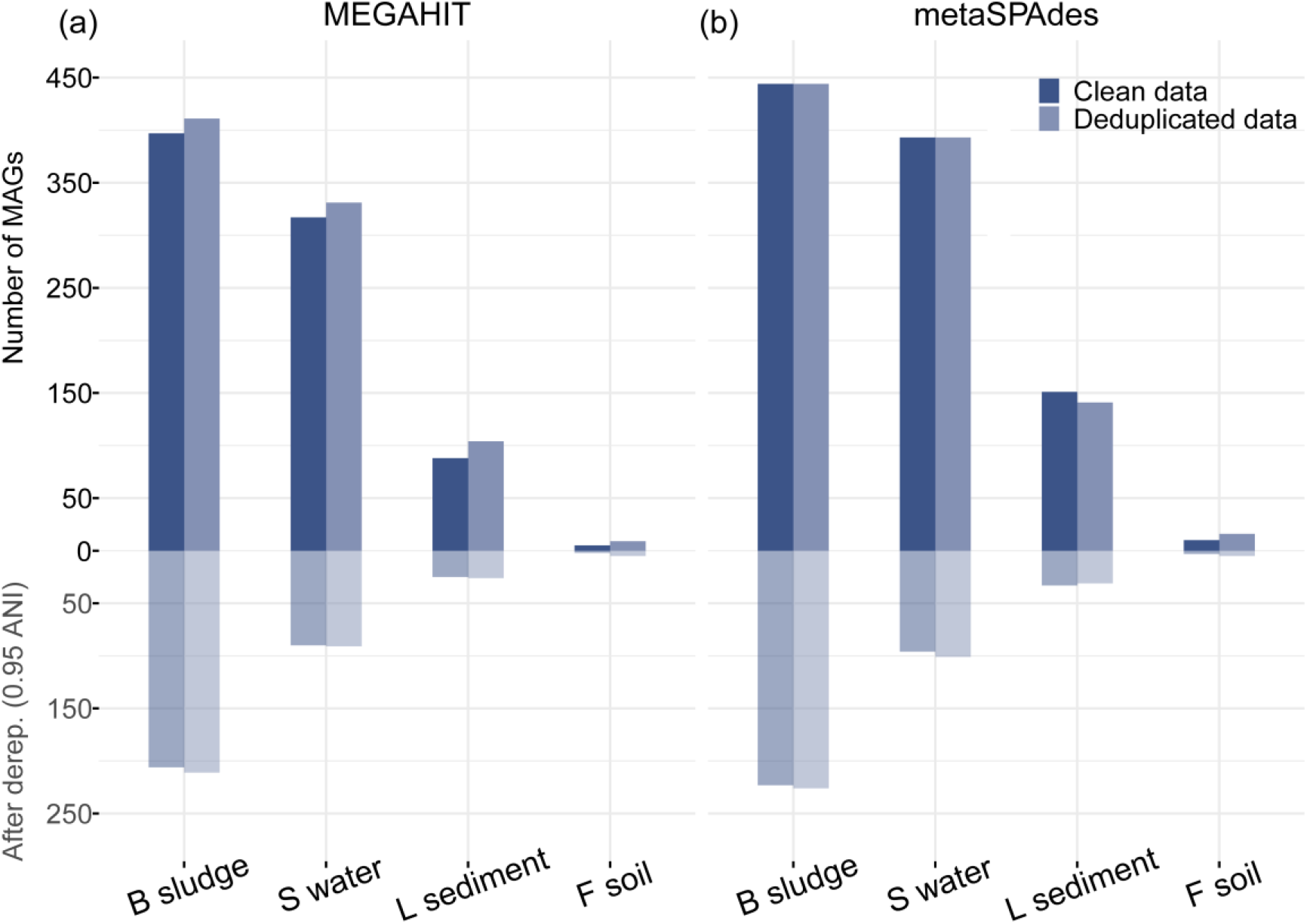
The number of MAGs recovered from MEGAHIT and metaSPAdes assemblies of clean and deduplicated data. The dereplication was conducted with cutoff of ANI > 95%. B sludge, S water, L sediment and F soil indicate bioreactor sludge, surface water, lake sediment and forest soil metagenomes.

**Table 1.**
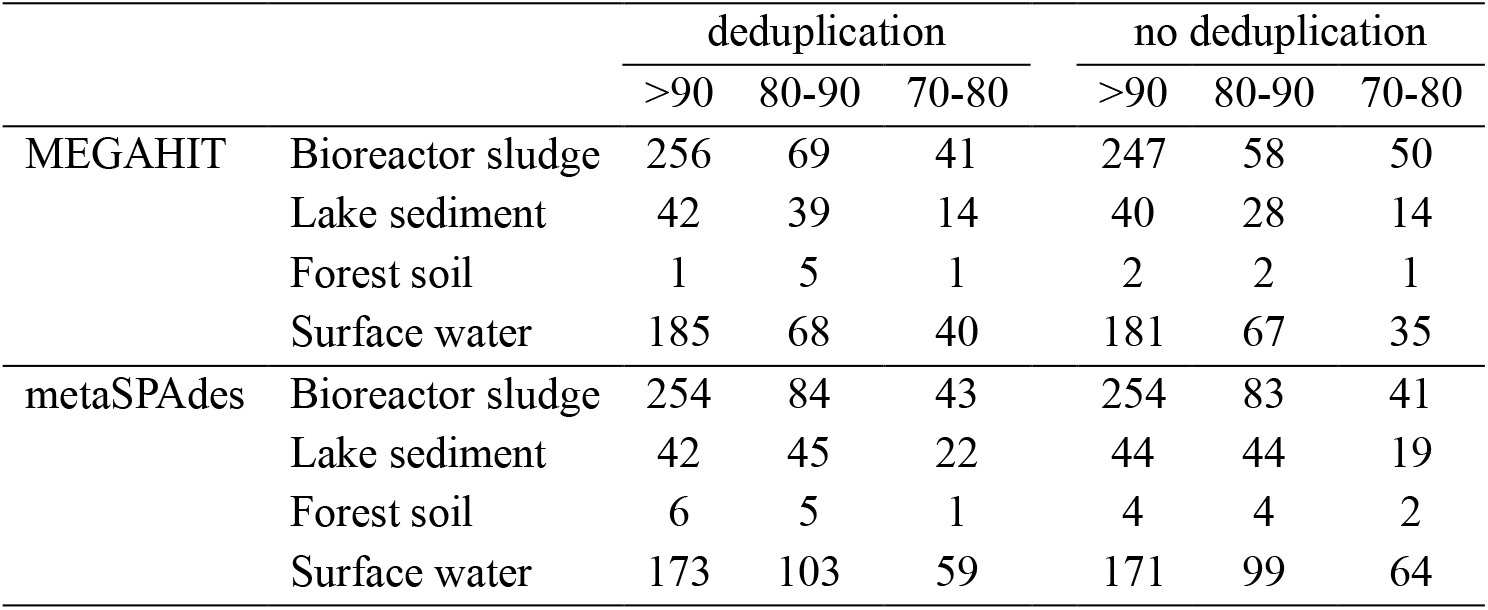
The number of high-quality MAGs

|  |  | deduplication |  |  | no deduplication |  |  |
| --- | --- | --- | --- | --- | --- | --- | --- |
|  |  | >90 | 80-90 | 70-80 | >90 | 80-90 | 70-80 |
| MEGAHIT | Bioreactor sludge | 256 | 69 | 41 | 247 | 58 | 50 |
|  | Lake sediment | 42 | 39 | 14 | 40 | 28 | 14 |
|  | Forest soil | 1 | 5 | 1 | 2 | 2 | 1 |
|  | Surface water | 185 | 68 | 40 | 181 | 67 | 35 |
| metaSPAdes | Bioreactor sludge | 254 | 84 | 43 | 254 | 83 | 41 |
|  | Lake sediment | 42 | 45 | 22 | 44 | 44 | 19 |
|  | Forest soil | 6 | 5 | 1 | 4 | 4 | 2 |
|  | Surface water | 173 | 103 | 59 | 171 | 99 | 64 |

The binning yields of metaSPAdes assemblies are remarkably higher than the binning yields of MEGAHIT assemblies, with 139 and 194 more MAGs recovered from deduplicated data and clean data, respectively (Fig. 3b). For details, the equivalent number of MAGs was recovered from deduplicated and clean data in bioreactor sludge (both 444 MAGs) and surface water (both 393 MAGs) metagenomes. For forest soil metagenomes, 6 more MAGs were recovered from metaSPAdes assemblies of deduplicated data, including 2 more MAGs with > 90% completeness and 1 more MAGs with > 80% completeness (Table 1). The number of high-quality MAGs (> 70% completeness) were slightly higher in deduplicated data (109 MAGs) than that in clean data (107 MAGs) of lake sediments (Table 1), although 10 more MAGs were recovered from clean data. Note that, after dereplication of MAGs (≥ 95% ANI), the number of species-level representative MAGs recovered from deduplicated data (363 MAGs) was higher than that recovered from clean data (355 MAGs). There were 226 versus 223, 101 versus 96, 31versus 33 and 5 versus 3 MAGs recovered from bioreactor sludge, surface water, lake sediment and forest soil datasets, respectively (Fig. 3b). Collectively, deduplication considerably promoted the binning yields of shot-gun metagenomic data on both total number of MAGs and number of species-level MAGs, particularly for the binning of MEGAHIT assemblies.

We also checked the chimera in MAGs through the taxonomic uniformity of 10 single-copy marker genes. Results showed that the disagreement rates of MAGs recovered from deduplicated data were lower than that of MAGs recovered from clean data in most of taxonomic level (Fig. S3). The total disagreement rate of MAGs recovered from MEGAHIT assemblies and metaSPAdes assemblies of clean data and deduplicated data were 40.1% versus 39.2% and 40.0% versus 38.6%, respectively. At genus level, the disagreement rates of MAGs recovered from metaSPAdes assemblies of deduplicated data (4.9%) were lower than that of clean data (6.1%) (Fig. S3b). These results indicates that less chimera present in MAGs recovered from deduplicated data compared with clean data.

In addition, we compared the above binning yields which based on their abundance correlation cross all samples from the same habitats (called cross-sample binning in this study) with another binning strategy, i.e., binning of assemblies based on their abundance profile in the corresponding one sample (called individual-sample binning in this study) which was adopted by several large-scale MAG reconstruction efforts (4,6,18). The results revealed that cross-sample binning remarkably recovered more MAGs than individual-sample binning from all four metagenomic datasets. A total of 855 and 630 MAGs were recovered by cross-sample binning and individual-sample binning of MEGAHIT assemblies of deduplicated metagenomes, respectively (Fig. S4a); and there were 994 and 761 in the across-sample binning and individual-sample binning of metaSPAdes assemblies (Fig. S4b). For details, 444 versus 354, 393 versus 314, 141 versus 79, and 16 versus 14 MAGs were recovered from cross-sample binning and individual-sample binning of metaSPAdes assemblies from deduplicated bioreactor sludge, surface water, lake sediment and forest soil metagenomes, respectively.

### Computational cost assessment of metagenomic assembly and binning

The access to affordable computational resources, e.g., maximum RAM memory and elapsed time, are critical important for bioinformatic analysis. Therefore, we evaluated the maximum RAM memory and elapsed time required for metagenomic assembly and binning of clean data and deduplicated data. The results showed that the deduplication significantly decreased the maximum memory requirement and time consumption during both MEGAHIT and metaSPAdes assembly (Fig. 4). For example, deduplication significantly decreased the memory requirements by 16.7% and 15.9% in the MEGAHIT assembly of bioreactor sludge and forest soil metagenomes on average, respectively (Fig. 4a). Meanwhile, deduplication significantly reduced elapsed time by 15.2%, 25.9%, 26.0% and 18.8% during MEGAHIT assembly of bioreactor sludge, surface water and lake sediment forest soil metagenomes, respectively (Fig. 4b). Likewise, the elapsed time was significantly reduced by 14.1%, 12.2%, 9.0% and 29.9% in the of metaSPAdes assembly of bioreactor sludge, surface water, lake sediment and forest soil metagenomes, respectively (Fig. 4d).

**Fig. 4.**
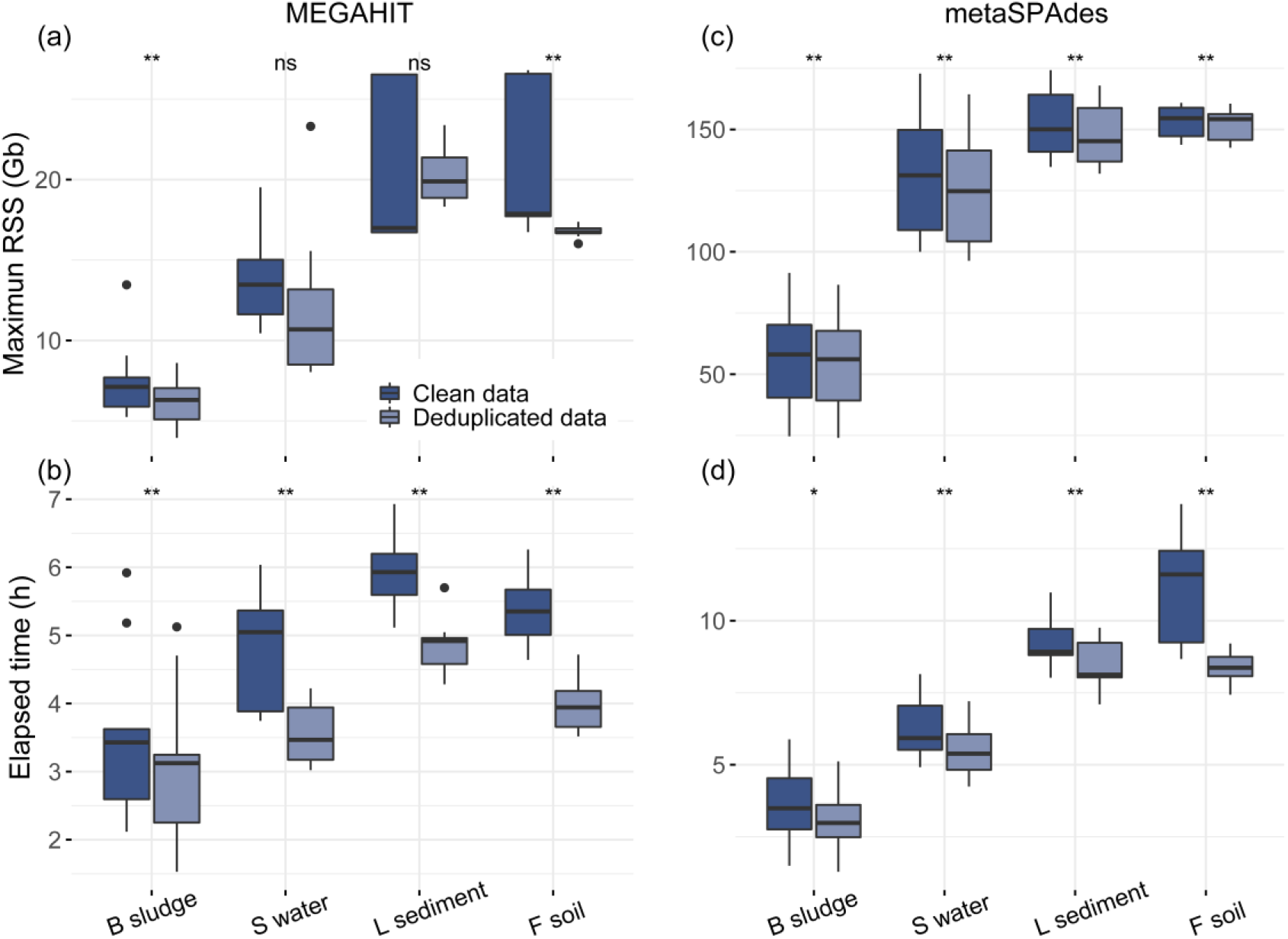
Time consumption and RAM memory requirement of assembly before and after deduplication of metagenomic data. Significance was checked using wilcoxon signed-rank test. “ns”, “*” and “**” indicate *p* > 0.05, *p* < 0.05 and *p* < 0.01, respectively.

Deduplication also reduced memory requirement and time duration of binning process (Fig. S5). Notably, deduplication significantly decreased the memory requirement by 13.7% and 5.9% in the binning of MEGAHIT and metaSPAdes assemblies of forest soil metagenomes, respectively. At the same times, the elapsed time was reduced by 25.3%, 19.3%, 28.6% and 21.4% in the binning of MEGAHIT assemblies of deduplicated bioreactor sludge, surface water, lake sediment, and forest soil metagenomes, respectively (Fig. S5b). Likewise, it was reduced by 15.6%, 11.2%, 13.2% and 15.0% in the binning of metaSPAdes assemblies of bioreactor sludge, surface water, lake sediment, and forest soil metagenomes, respectively (Fig. S5d). In summary, deduplication considerably saved elapsed time and RAM memory in most of the studied assembly and binning cases.

## Discussion

Over the last decade, the improvement of next-generation metagenomic sequencing technologies and advances in computational methods have revolutionized the field of microbiology and microbial ecology (18), which enables the genome-centric studies through the recovery of metagenome-assembled genomes (MAGs) from various complex environments, including the global ocean (19), cow rumen (20), human gut (21,22), aquifer (23) and soil (24). MAGs recovered from metagenomic data allowed robust and detailed qualitative views of functional and metabolic aspects of microbiome within a genomic context with high taxonomic and phylogenetic resolution (2,3). Further, the genome reconstruction of uncultivated bacteria and archaea has substantially expanded and populated the microbial tree of life (4,25,26), and yielded significant insights into evolutionary relationships and metabolic properties of uncultivated fraction of microbiomes (23). However, the presence of duplicate reads (derived from the library construction of metagenomic sequencing) in metagenomic data and their influence on metagenomic assembly and binning have been overlooked for a long time.

Therefore, in this study, we systematically investigated the impacts of duplicate reads on assembly and binning of real metagenomic datasets with various microbiome complexity, i.e., bioreactor sludge, surface water, lake sediment and forest soil. The results showed that deduplication improved the construction of long contigs, and the size of longest contig for most of samples were considerably improved after deduplication, particularly for metaSPAdes assembly (Fig. 2). The improvement of contig length might contribute to the removal of duplicate reads in metagenomic data, which reduced the coverage bias of kmer, leading to continuous assembly of contigs (27,28). Long contigs could enable a more sensitive detection of large complex genomic features such as clustered regularly interspaced short palindromic repeats (CRISPR) (29), polyketide synthase (PKS) or non-ribosomal peptide synthase (NRPS) gene clusters encoding for secondary metabolites (24).

Furthermore, deduplication is found to considerably improve the binning yields. For example, the binning of MEGAHIT assemblies from deduplicated data yields more MAGs than that from clean data of bioreactor sludge, surface water, lake sediment and forest soil metagenomes, and deduplication also increases the number of species-level representative MAGs recovered from both MEGAHIT and metaSPAdes assemblies (Fig. 3). Based on previous studies, contig coverage abundance is an important metric in the widely used binning tools (30). Therefore, the presence of duplicate reads might affect coverage abundance of contigs, leading to the increase of contamination level of MAGs. Our results also confirmed the decrease of contamination level of MAGs recovered from deduplicated data, as the results showed that deduplication decreased the disagreement rates of taxonomic annotations of MAGs (Fig. S3). In addition, across-sample binning strategy is recommended in the recovery of MAGs from metagenomes, as more MAGs were recovered in this study comparing with individual-sample binning (Fig. S4). In the across-sample binning, each assembly has the coverage information of all samples from the same habitat. Therefore, this might contribute to more efficient and accurate binning of assemblies, as also indicated by binning developers(31).

Besides improving metagenomic assembly and binning, deduplication is also demonstrated to greatly decrease computational cost (i.e., elapsed time and RAM memory) of assembly and binning (Fig. 4 and Fig. S5), because of the reduction of duplicate reads (by 10%-20% in this study, Dataset S1). Although metagenomes from different environmental habitats have similar data size in this study, the demand of computational resources was greatly distinct among samples, depending on their microbiome diversity and complexity. The higher microbiome diversity the samples (e.g., lake sediment and forest soil) have, the longer elapsed time and higher memory requirement will be taken to complete the assembly and binning. Therefore, the assembly of big-size complex metagenome will greatly challenge the computational capacity, so it is advisable to remove duplicate reads in order to decrease demand of computational resources. Collectively, it is demonstrated that deduplication improves both the binning yields and cost-efficiency for all four groups of metagenomes, i.e., bioreactor sludge, surface water, lake sediment and forest soil, tested in this study. Therefore, it is recommended to remove duplicate reads before metagenomic assembly and binning.

However, it should not be neglected that deduplication may remove the natural duplicate reads from predominant species of an uneven microbiota, even though the probability of such an event in metagenomic data of environmental samples is extremely low (10). Natural duplicate reads might increase in less complex microbiota samples (32) such as enriched bioreactor sludge and environmental microbial cultures, so specific technologies (e.g, Unique Molecular Identifiers), which give unique label to each insert sequence (33), can be applied to inspect natural duplicates and accurately remove artificial ones when desired. The artificial duplicate reads do not represent any biologically meaningful information, and cause the unnecessary waste of sequencing and computational resources. Therefore, it is better to avoid or minimize the generation of such duplicate reads from its source of production (i.e., metagenomic library preparation). For example, providing enough genomic DNA is alwayes recommended for sequencing library construction to minimize PCR cycles, as it was reported that PCR duplicates arise from a lack of DNA complexity (unique templates) due to low levels or quality of input DNA (32,34). Another way to aviod the generation of artifcial duplicate reads is to develop methods of library constrction without PCR amplification (9,28), which can aviod artificial duplicate reads and amplification bias from the source.

## Materials and methods

### Sample collection and metagenomic sequencing

Environmental samples were collected from four engineered or natural microbial ecosystems, including nine activated sludge samples of a bioreactor run in our laboratory at the Westlake University (Hangzhou, China), nine surface water samples of salt marsh in Dafeng Milu national nature preserves (Yancheng, China), nine sediment samples from Taihu lake (Wuxi, China), and nine soil samples from Maolan karst forest (Libo, China). Total genomic DNA of each sample was extracted using FastDNA Spin Kit for Soil (MP Biomedicals, USA) following the manufacturer’s instructions, and was sequenced on the Illumina NovaSeq 6000 platform with a paired-end 150 bp strategy at Novogene Corporation (Beijing, China). Each sample has a data size of ~ 10 Gbp, and more details on the metagenomes used could be found in Dataset S1.

### Metagenomic data preprocessing, assembly, and binning

Raw reads were filtered to remove low quality reads, N containing reads, and adapters using fastp (v0.23.1) (35), yielding the clean data. Then, duplicate reads in clean data were removed using fastp (v0.23.1), generating the deduplicated data (See detailed information in Dataset S1). In order to evaluate the impacts of duplicate reads on metagenomic assembly and binning, the clean-read datasets and deduplication datasets were individually *de novo* assembled using both MEGAHIT (v1.2.9) (36) and metaSPAdes (v3.14.1) (27), with default parameters. The resulting assembly of each sample was subsequently clustered into genome bins. Briefly, the coverage profiles of each assembly were calculated by mapping reads of every sample from the same habitat, which means that each assembly has the coverage information of nine samples in this study. The coverage profiles of each assembly were then used to inform automated binning of MetaBAT2 (v2.12.1) (37), which was called cross-sample binning in this study. We also compared the above binning yields with the yields of another binning strategy, called individual-sample binning in this study, in which the coverage profile of each assembly was generated by only mapping clean reads of the corresponding one sample, and was used to inform binning as the above described.

### Results assessment, statistical analyses, and visualization

The microbiome complexity of each metagenome was assessed using Nonpareil (v3.4.1) (38), a statistical program that uses read redundancy to estimate sequence coverage. Metagenomic assembly of each sample was evaluated using QUAST (v5.0.2) (39). The quality of metagenome-assembled genomes (MAGs) was assessed using checkM (v1.2.0) (40), and the MAGs with an overall quality ≥50% (completeness–5×contamination) were considered as eligible bins. Species-level representative MAGs were obtained by dereplicating the above MAGs using dRep (v3.0.0) (41). The average nucleotide identity (ANI) was calculated using FastANI (v1.33) (42). The chimera in MAGs was evaluated through taxonomic uniformity using STAG (v0.8.2, https://github.com/zellerlab/stag). We annotated 10 single-copy marker genes and evaluated the homogeneity of their taxonomic annotation for each MAG, as follows: “No annotation” if less than two marker genes were annotated; “Agreeing” if all marker genes had the same annotation; and “Disagreeing” if different annotation of marker genes presented. The disagreement rate was calculated by the number of “Disagreeing” MAGs divided by the number of all MAGs excluding “No annotation”. Elapsed time and RAM memory taken by software to complete the metagenomic assembly and binning were recorded using an in-house bash script. R (v4.0.4) was used to perform statistical analyses and figure plotting.

## Acknowledgement

This work was supported by Zhejiang Provincial Natural Science Foundation of China under Grant No. LR22D010001 and National Natural Science Foundation of China under Grant No. 51908467. We thank Ms. Yisong Xu for laboratory management supports and Mr. Guoqing Zhang for laboratory server maintenance and meaningful discussion. We acknowledge the Research Center for Industries of the Future (RCIF) at Westlake University for supporting this work, and thank the Westlake University HPC Center for computation support.

## Data Availability

The metagenomic sequencing data used for methodology evaluation in this study are contributed from unpublished projects of lab members under the accession number of CNP0003575 in the China National GeneBank database.

